# Altered sleep intensity upon DBS to hypothalamic sleep-wake centers in rats

**DOI:** 10.1101/2021.08.11.455908

**Authors:** S. Masneuf, L.L. Imbach, F. Büchele, G. Colacicco, M. Penner, C.G. Moreira, C. Ineichen, A. Jahanshahi, Y. Temel, C.R. Baumann, D. Noain

## Abstract

Deep brain stimulation (DBS) has been scarcely investigated in the field of sleep research. We hypothesize that DBS onto hypothalamic sleep- and wake-promoting centers will produce significant neuromodulatory effects, and potentially become a therapeutic strategy for patients suffering severe, drug-refractory sleep-wake disturbances. We aimed to investigate whether continuous electrical high-frequency DBS, such as that often implemented in clinical practice, in the ventrolateral preoptic nucleus (VLPO) or the perifornical area of the posterior lateral hypothalamus (PeFLH), significantly modulates sleep-wake characteristics and behavior. We implanted healthy rats with electroencephalographic/electromyographic electrodes and recorded vigilance states in parallel to bilateral bipolar stimulation of VLPO and PeFLH at 125 Hz at 90 μA over 24 h to test the modulating effects of DBS on sleep-wake proportions, stability and spectral power in relation to baseline. We unexpectedly found that VLPO DBS at 125 Hz deepens slow-wave sleep as measured by increased delta power, while sleep proportions and fragmentation remain unaffected. Thus, the intensity, but not the amount of sleep or its stability, is modulated. Similarly, the proportion and stability of vigilance states remained altogether unaltered upon PeFLH DBS but, in contrast to VLPO, 125 Hz stimulation unexpectedly weakened SWS, evidenced by reduced delta power. This study provides novel insights into non-acute functional outputs of major sleep-wake centers in the rat brain in response to electrical high-frequency stimulation, a paradigm frequently used in human DBS. In the conditions assayed, while exerting no major effects on sleep-wake architecture, hypothalamic high-frequency stimulation arises as a provocative sleep intensity-modulating approach.

## Introduction

The regulation of physiological sleep and wakefulness relies on the equilibrium of a well-explored network of wake- and sleep-promoting centers in the brain. Mutual inhibition between these centers has been proposed as main regulatory mechanism in the flip-flop switch model [1, 2], where effective sleep requires the suppression of arousal systems by the mainly inhibitory GABAergic ventrolateral preoptic nucleus (VLPO) in the anterior hypothalamus [3, 4], whose activation via agonism of adenosine receptors increases non-rapid eye movement (NREM) sleep [5, 6]. Additional inhibition of wake-promoting neurons by adenosine builds up on this neurotransmitter’s sleep-promoting action [7]. As a counterpart, the perifornical area of the posterior lateral hypothalamus (PeFLH) is densely populated, among other cell populations, by hypocretin (orexin) neurons, implicated in the facilitation of arousal [8]. The PeFLH region may also be an important organizer of the normal succession of stages during the sleep-wake cycle through excitatory reciprocal feedback to monoaminergic centers including the locus coeruleus (LC), a major arousal hub [8].

The complex regulation of sleep and wakefulness by counteracting and heterogeneous hypothalamic and brainstem nuclei, including VLPO and PeFLH, has been demonstrated in the last decades using stereotaxic injections of toxins/receptor agonists, as well as by opto- and pharmacogenetics [9–12]. Such approaches have been very useful and successful in specifically targeting distinct neuronal populations within heterogeneous brain areas and modulating sleep phenotypes in animals, thus identifying the basic nature of the wake- and sleep-controlling nuclei. However, while precise information is obtained from these approaches when they are acutely applied, their neuromodulatory effects during long-term (hours, days) applications have not been characterized. Additionally, this molecular approach presents some practical limitations towards cell types and species [13], with restricted genetic tools and inaccessibility to larger brains [14], which joins other factors such as the irreversibility of the genetic manipulation and the unknown long-term consequences of viral expression, and potential protein accumulation, as caveats for eventual clinical applications.

Overall, clinically accessible approaches, such as electrical deep brain stimulation (DBS), have to be explored to determine whether targeting sleep-wake-controlling brain areas is a valuable therapeutic strategy offering significant neuromodulatory effects.

Here we aimed to study the effects of high frequency electrical neuromodulation of the VLPO and PeFLH nuclei, as representative sleep-wake-controlling hypothalamic targets, to determine whether these regions can serve as valuable putative therapeutic targets in further animal and human studies on the treatment of severe, drug-refractory sleep-wake disturbances. We thus tested the effect of continuous electrical high frequency stimulation (HFS, often implemented in clinical practice for reversible functional ablation of the targeted nuclei; [15]) on sleep-wake behavior, stability, and intensity as assessed by electroencephalographic/electromyographic (EEG/EMG) recordings in light, dark and 24 h periods. We hypothesize that HFS of the VLPO area will decrease NREM sleep proportion, stability, and/or intensity - by functionally inhibiting its sleep-promoting action -, whereas HFS of the PeFLH region will decrease wakefulness proportion, stability and/or intensity - by functionally disabling the arousal/wake-promoting nucleus’ function -.

## Materials and methods

### Experimental design

A total of 14 animals were chronically instrumented with EEG/EMG headsets and DBS leads into the VLPO, from which 5 animals were excluded from the analysis due to histological determination of mistargeting of the DBS leads, leaving 9 available animals. In those 9 animals, we performed sleep-wake proportions, fragmentation and delta power analyses upon HFS in light, dark and 24 h periods. For each analysis, statistical outliers, defined as cases with behavioral scores > 2 standard deviations from each overall group mean, and/or animals presenting technical issues with the EEG/EMG, were additionally excluded from the analysis. Thus, for the VLPO, the remaining number of animals analyzed was n=7 during the light period (1 statistical outlier and 1 technical failure in entire EEG) and n=6 during the dark period (1 statistical outlier, 1 technical failure in entire EEG, and 1 technical issue during the dark period) for all parameters.

A total of 8 animals were chronically instrumented with EEG/EMG headsets and DBS leads into the PeFLH, from which 1 animal was excluded from the analysis due to histological determination of DBS leads mistargeting. In the remaining 7 animals, we analyzed sleep-wake proportions, fragmentation and delta power analyses also in light and dark periods and per 24 h. No statistical outliers were found in the analyses, whereas again we excluded one animal presenting technical issues in the entire EEG/EMG. Thus, in the PeFLH, the remaining number of animals analyzed was n=6 for all parameters.

### Animals

We included adult male Sprague-Dawley rats (Charles River Laboratories International Inc, Germany) weighing between 290-380 g at the time of surgery. The rats were housed individually on a 12:12 light:dark cycle with food and water available *ad libitum* throughout the experiments. In compliance with ethical regulations and to explore potential side effects that would cause eventual protocol discontinuation, we monitored animals’ weight daily while in EEG/EMG recordings/DBS sessions as well as home-cage activity after interventions. We additionally checked body temperature before and after the interventions. The animal room temperature was constantly maintained at 21-24°C. All experiments were approved by the veterinary office of the canton of Zurich and conducted according to the local guidelines for care and use of laboratory animals under license ZH205/12.

### Surgical procedures for EEG/EMG and DBS electrode implantation

We anesthetized rats with isoflurane (4.5% for induction, 2.5% for maintenance) and injected them with buprenorphine (0.05 mg/kg, s.c.) for analgesia. We monitored wound healing, body weight and home-cage activity of the animals on a daily basis over the first week after surgeries and weekly thereafter.

We performed the EEG/EMG and DBS implantation procedures using adapted versions from previous established protocols [16, 17]. Briefly, we positioned the animals in a standard stereotactic apparatus (model 1900, Kopf Instruments, Tujunga, CA, USA) over a temperature-controlled pad, and made a midline incision exposing the skull. We made burr holes over the position of DBS coordinates followed by duratomy. Before insertion of the DBS electrodes, we frontally placed two anchoring screws (M1, one per hemisphere) and inserted four screws for peridural EEG recording, one pair per hemisphere over the parietal cortex. Additionally, we inserted a pair of gold wires into the rats’ neck muscles, which served as EMG electrodes for monitoring muscle tone. All EEG/EMG electrodes were connected to a plug by soldering to stainless steel wires.

We implanted DBS electrodes on targets after adjustment of the initial coordinates: (i) VLPO (AP −0.04 mm, ML 0.8 mm, DV −10 mm) using a factor obtained by dividing the measured individual Bregma-Lambda distance by the reference measure from the Atlas [18], (ii) PeFLH (AP −2.9 mm, ML 1 mm; DV −9 mm) with AP −2.7 mm applied for measured Bregma-Lambda distance < 6.9 mm and AP −3.1 mm applied for measured Bregma-Lambda distance ≥ 6.9 mm. Finally, we cemented all headpieces to the skull as illustrated in **Fig 1**.

**Fig 1.**
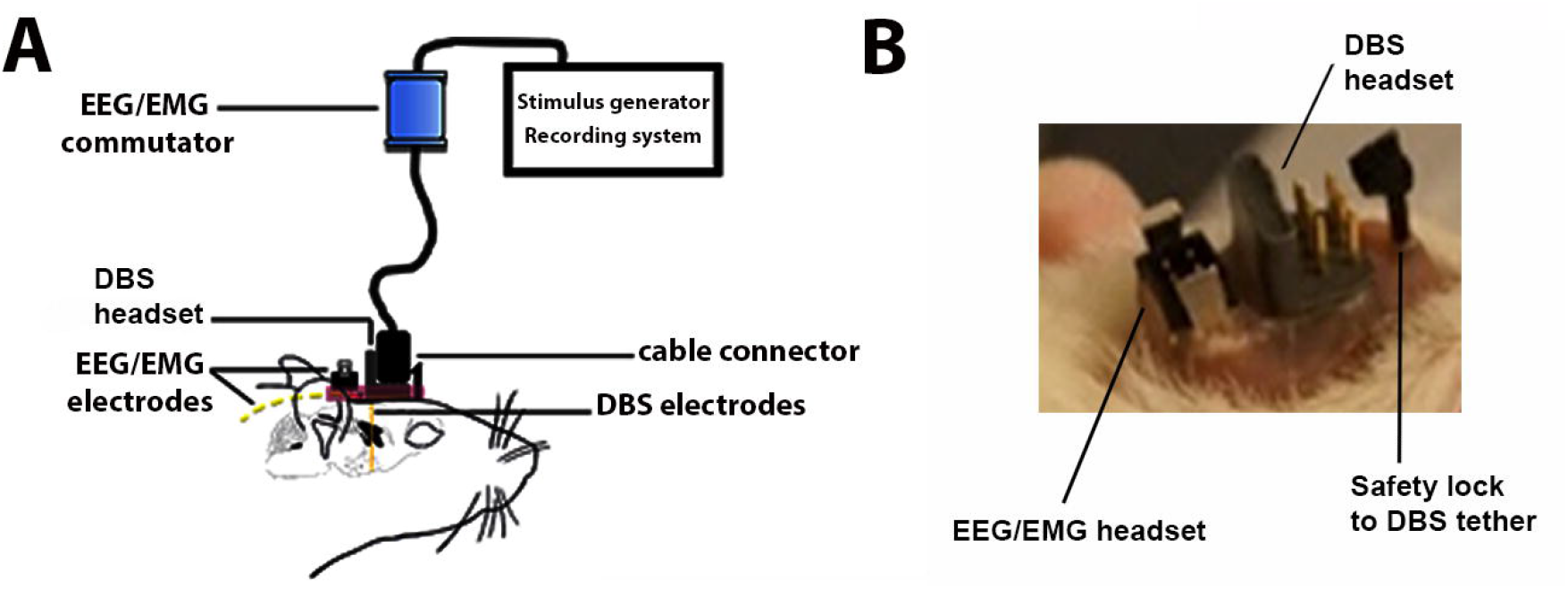
DBS and EEG/EMG electrode implantation in rats. (**A**) Scheme of the experimental set up. The animal is freely moving in its cage while connected to a swivel cable, which allows both stimulation of the targeted brain structure and recording of the sleep-wake patterns. (**B**) Post-surgical DBS-EEG/EMG electrode construction.

### Deep brain stimulation: hardware and characteristics

In our experimental set up, we used bilateral concentric bipolar DBS electrodes as previously described [16], stereotactically implanted into the VLPO and PeFLH. The gold-plated electrodes, composed of an inner platinum-iridium wire, functioning as the negative pole, and an outer stainless steel layer as the positive one, produce a concentrated current field around the tip of the electrode. The maximum outer diameter of the electrode is about 250 μm with a tip diameter of approximatively 50 μm (Technomed, Beek, The Netherlands).

We used a stimulus of rectangular shape in a current-controlled paradigm and applied a bipolar biphasic (60 μs negative - 60 μs positive) stimulation, which bears the advantage of potential translation into clinical studies.

### Selected DBS parameters

Based on the most commonly used parameter in the literature, we selected 125 Hz as frequency of stimulation, fixing the negative pulse width at 60 μs as regularly used in clinical studies. We based the choice of the pulse amplitude on a priori simulations aiming at investigating the distribution of the electrical field in brain tissue [19]. We modeled the electrode position in the VLPO, as proxy structure due to its relatively small and well-defined anatomy. For this, we used the final outcome of the finite element method of modeling and simulation, a widely applied numerical technique for calculating approximate solutions of general partial differential equations [19]. The model integrated the electrode configuration (top *vs*. center) and aspects of the local structure of the surrounding region. We split the region of interest into 4 quadrants and neglected overlapping effects of the resulting borders. We applied a physics controlled mesh, and set the density with an application of a spatial resolution of 1 μm for the region close to the electrode (0 to 0.15 mm). For the distal region (0.15 mm to 10 mm), we used a spatial resolution of 10 μm. We set tissue conductivity to 0.07 S/m (within common frequency ranges), at body temperature and of bovine origin due to lack of data for the rat [20] and the pulse width to 0.06 ms. Moreover, we accommodated for dielectric tissue properties represented by optic chiasm −och− (medial to the left VLPO) and median forebrain bundle −mfbb− (lateral to the left VLPO). We assessed the distance of mfbb and och by taking the average of two brain slices at AP level 0.00 and −0.12 mm from bregma [18] where we preferentially targeted the VLPO. We performed simulations at 40, 90 and 150 μA and observed no summation effects (i.e. absence of rest-energy when the second pulse is applied). Accordingly, we neglected effects of frequency. To increase the comparability between the different simulations using either 40, 90 or 150 μA, the absolute field strength (3.75 V/mm) was kept constant. The simulations were based on relative (and not absolute) electrical field strengths. The final outcome of the simulations revealed distributions of the electrical fields beyond the target-region at the three intensities of interest for both top and center configurations. Stimulation at 90 and 150 μA would target 50 to 100% of VLPO volume, depending on the position of the electrodes (i.e. top or center). In comparison to 90 μA, however, at 150 μA the boundaries of the main surrounding structures (och and mfbb) would largely be affected by the stimulation. Still, even at lower intensities of stimulation, the modeling results indicate that we cannot avoid affecting, at different degrees, the surrounding structures of VLPO, predicting that targeting a specific region without any current leakage is unlikely. Thus, we chose to stimulate at 90 μA to minimize the main leakage effects while substantially targeting the VLPO. We also chose this amplitude of stimulation for the PeFLH area, a relatively bigger area than VLPO, assuming we will be further limiting leakage effects.

### Experimental protocols and setup

We single-housed the rats following surgery and granted a recovery of at least 2 weeks to all animals before further interventions. We connected the DBS electrodes to a stimulation device (model STG4008-1.6mA, Multi Channel Systems MCS GmbH, Reutlingen, Germany) in parallel to EEG/EMG electrodes through externalized cables that hang from a rotating swivel at the top of the cage, allowing free motion of the animals inside the experimental cage (**Fig 1**). We monitored stimulus delivery using an oscilloscope. Following a setup adaptation period of one to two days, we conducted EEG/EMG recordings for 24 h during two consecutive days, one before (DBS OFF) and one during (DBS ON) continuous bilateral electrical stimulation (**Fig 2A**).

**Fig 2.**
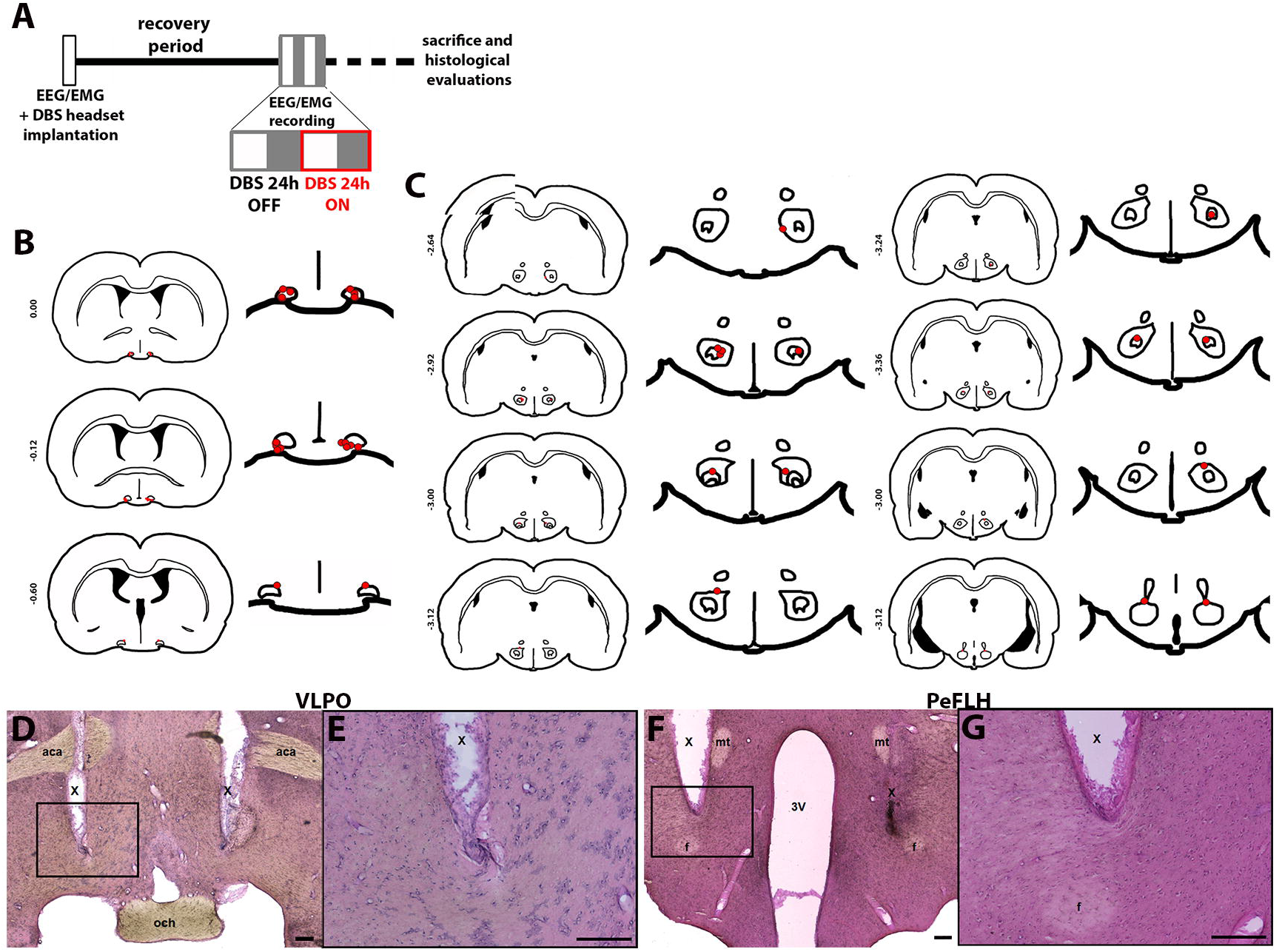
Experimental design and DBS electrodes placement. (**A**) Rats were implanted with EEG/EMG electrodes and DBS headset for the recording of vigilance states and stimulation respectively, followed by a recovery period of minimum two weeks. After one to two days of adaptation, EEG/EMG recordings were performed for two consecutive days: before (DBS OFF, 24 h) and during (DBS ON, 24 h) stimulation. Animals were then sacrificed, and electrode location and tissue integrity were verified through hematoxylin-eosin staining analyses. White squares: light period; grey squares: dark period. (**B**) and (**C**) Schematic coronal sections indicating site of electrode placements in VLPO (n=9) (**B**) and PeFLH (n=7) (**C**) regions (Bregma coordinates in mm) with target outlined in black and electrode tip pointed in red. (**D-G**) Representative low and high-magnification micrographs illustrating electrode targeting of VLPO (**D**, **E**) and PeFLH (**F**, **G**). Scale bars represent 200 μm in all pictures. VLPO: ventrolateral preoptic area; PeFLH: perifornical lateral hypothalamic area; X: electrode track; aca: anterior commissure, anterior part; och: optic chiasm; f: fornix, 3V: third ventricle; mt: mamillo-thalamic tract.

### EEG/EMG recording, scoring and analysis

We sampled EEG and EMG at 200 Hz and amplified and processed the signals by an analog-to-digital converter. We used EMBLA hardware and Somnologica-3 software (Medcare Flaga). We discarded activity in the 50 Hz band from the analysis because of power line artifacts. We obtained power spectra of the EEG by discrete Fourier transformation (range: 0.5-100 Hz; frequency resolution: 0.25 Hz; time resolution: consecutive 4 s epochs; window function: Hanning).

We performed blinded visual scoring of acquired EEG manually as described previously [17]. We excluded artifacts by visual review of the raw data and identified and scored three vigilance states based on EEG/EMG patterns: wakefulness (WAKE), NREM sleep, and rapid eye movement (REM) sleep. We divided the 24 h scoring sessions into light and dark periods or into 1 h intervals, and assessed the proportion of the vigilance states separately for each period. To provide a quantitative measure of sleep fragmentation, we calculated the sleep fragmentation index as follows: a behavioral state bout was defined as a consecutive series of epochs in the same behavioral state without state transitions. The resulting amount of behavioral state bouts was then divided by the total number of 4 s epochs in the same sleep stage, resulting in a comparable measure for fragmentation between 0 and 1 [17, 21]. We also determined the total delta power of NREM sleep, calculated as the summarized power in the slow-wave activity (SWA) band (0.5-4 Hz), before and during stimulation. We performed all signal processing and analyses as described using MATLAB (MathWorks).

Furthermore, we analyzed the build-up of delta power (relative delta power) upon transition into consolidated NREM sleep as described earlier [22]. To this end, we identified all consolidated NREM episodes lasting longer than 6 minutes. This value was determined based on established criteria (7 minutes; [23]) and adapted empirically to ensure a significant amount of consolidated sleep bouts in the analysis (6 minutes time frame).

### Electrode placement confirmation

Upon completion of the experiments, we sacrificed the rats via intracardiac perfusion as previously described [24]. We verified correct electrode placement and visualized potential tissue damage due to the stimulation by hematoxylin-eosin stainings in coronal 40 μm fixed brain sections (**Fig 2B-G**) [18].

### Statistical analyses

We expressed the light, dark and 24 h proportions of vigilance states, and the logarithm of delta power as medians and quartiles with 95% confidence intervals (CI), while fragmentation and data presented in 1 h intervals were plotted as means ± S.E.M. The full spectrum EEG data were reported as absolute values. Bivariate comparison of delta power and sleep fragmentation ON *vs*. OFF stimulation was done by Wilcoxon’s signed rank tests and Bonferroni corrections (n=4 tests). A two-way analysis of variance (ANOVA) with repeated measures (factor band - 4 levels: alpha, beta/gamma, delta, theta; factor DBS - 2 levels: OFF, ON) was used for the statistical assessment of the full spectrum EEG data for the three vigilance states and for the analysis of delta-build up in time, followed by Bonferroni post-hoc tests as appropriate. One-way ANOVA with repeated measures (factor: hour - 24 levels: 12 hours OFF *vs*. ON) was additionally applied within each light and dark periods, for the detailed analysis of the time course of the vigilance states and delta power, followed by Student-Newman-Keuls post-hoc tests as appropriate. We used a threshold for statistical significance of P ≤ 0.05. We performed all statistical analyses using StatView^®^ (SAS Institute Inc., USA) and R (Team).

## Results

### No distinct side effects or tissue damage upon DBS

Importantly, we did not observe side effects in respect to body weight, body temperature, and/or overall home cage behavior (data not shown) in association with our interventions. Evaluation of hematoxylin-eosin staining revealed no significant tissue damage, besides the electrode tracks, in or around the target structures (**Fig 2**).

### DBS modulation effect on sleep-wake behavior and stability

Detailed analysis of the 24 h time course of the vigilance states in 1 h data intervals did not reveal specific time windows of neither VLPO (**Fig 3A**) nor PeFLH (**Fig 3B**) DBS effects for WAKE (VLPO, light period: F(23,138) = 4.46, P < 0.0001; dark period: F(23, 115) = 2.72, P = 0.0002; followed by non-significant Student-Newman-Keuls post-hoc comparisons; PeFLH, light period: F(23, 115) = 2.26, P = 0.0025; dark period: F(23, 115) = 4.61, P < 0.0001; followed by non-significant Student-Newman-Keuls post-hoc comparisons), NREM sleep (VLPO, light period: F(23,138) = 3.79, P < 0.0001; dark period: F(23,115) = 2.65, P = 0.0004; followed by non-significant Student-Newman-Keuls post-hoc comparisons; PeFLH, light period: F(23, 115) = 2.11, P = 0.0052; dark period: F(23, 115) = 4.16, P < 0.0001; followed by non-significant Student-Newman-Keuls post-hoc comparisons), and REM sleep (VLPO, light period: F(23,138) = 3.79, P < 0.0001; dark period: F(23,115) = 2.26, P = 0.0026; followed by non-significant Student-Newman-Keuls post-hoc comparisons; PeFLH, light period: F(23, 115) = 2.28, P = 0.0023; dark period: F(23, 115) = 4.71, P < 0.0001; followed by non-significant Student-Newman-Keuls post-hoc comparisons). We additionally corroborated no significant changes in light, dark and per 24 h sleep-wake proportions upon HFS in either VLPO (**Fig 3C**) or PeFLH (**Fig 3D**). Stability of sleep-wake behavior - as calculated by the index of fragmentation in both dark and light periods for all three vigilance states - was also unaltered upon VLPO (**Fig 4A**) and PeFLH (**Fig 4B**) HFS.

**Fig 3.**
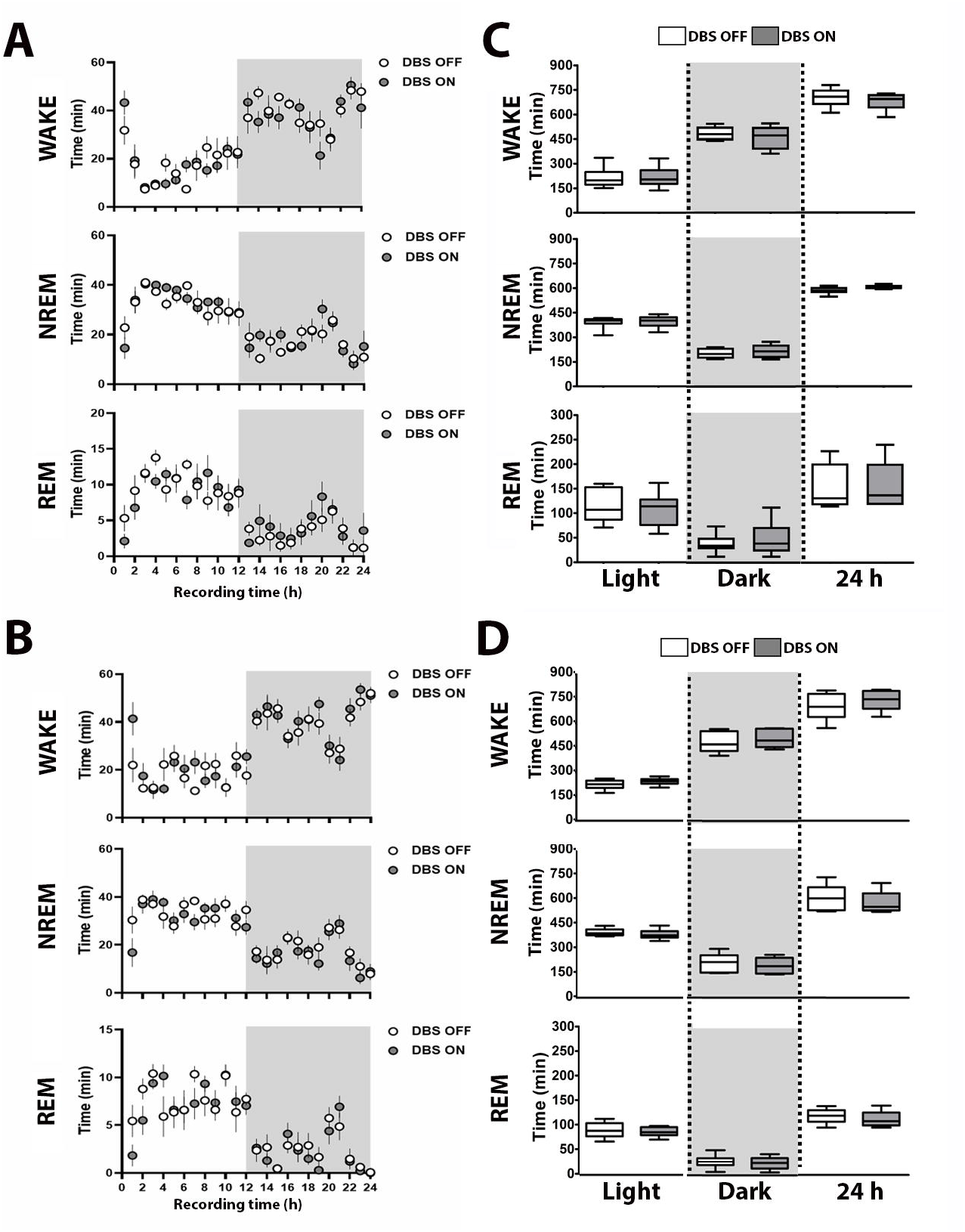
Effect of VLPO and PeFLH HFS on vigilance state proportions. (**A-B**) No changes in the hourly proportions of vigilance states nor in the (**C-D**) light, dark or total 24 h periods, when comparing data before (DBS OFF) and during (DBS ON) 125 Hz stimulation of VLPO (**A-C**) and PeFLH (**B-D**). Vigilance state proportions data are expressed as medians and quartiles with 95% CI. Data in 1 h intervals are means ± S.E.M. Wilcoxon’s signed rank tests and Bonferroni corrections. One-way repeated measures ANOVA followed by Student-Newman-Keuls tests. n=6-7 per group. Two statistical outliers (criteria: scores > 2 standard deviations; 1 for the analysis of the light period and 1 for the analysis of the dark period) were excluded from the analysis. WAKE: wakefulness, NREM: non-rapid eye movement sleep, REM: rapid eye movement sleep; min: minutes.

**Fig 4.**
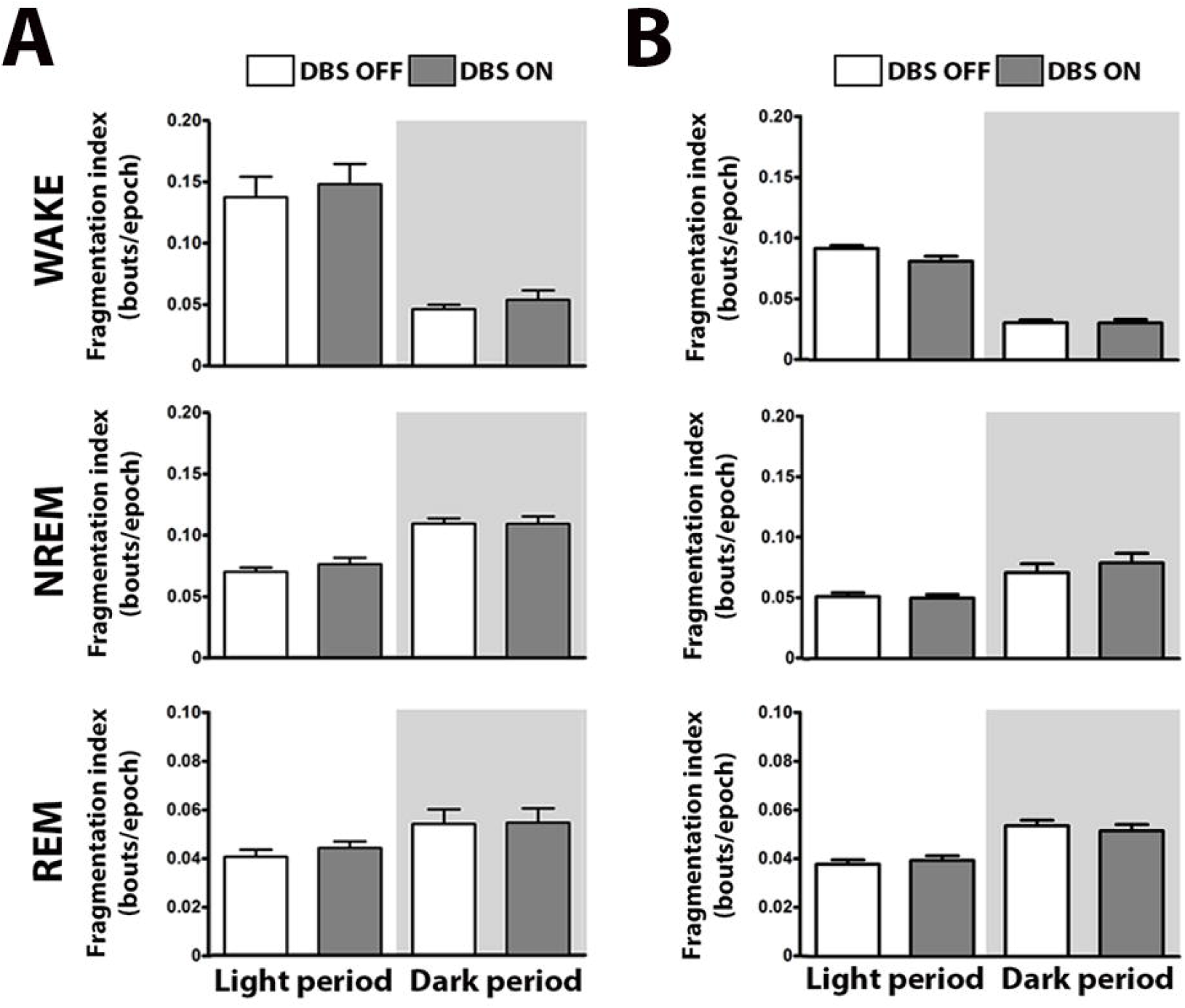
Effect of VLPO and PeFLH HFS on sleep fragmentation index. Behavioral state stability was maintained as represented by the absence of changes in the fragmentation index, when comparing data before (DBS OFF) and during (DBS ON) stimulation of VLPO (**A**) and PeFLH (**B**). Fragmentation index data are expressed as means ± S.E.M. Wilcoxon’s signed rank tests and Bonferroni corrections. n=6-7 per group. Two statistical outliers (criteria: scores > 2 standard deviations; 1 for the analysis of the light period and 1 for the analysis of the dark period) were excluded from the analysis. WAKE: wakefulness, NREM: non-rapid eye movement sleep, REM: rapid eye movement sleep.

### DBS modulation effect on sleep intensity

Following no changes in sleep-wake behavioral patterns and along the hypothesis that DBS could alternatively have specifically modulated intensity of sleep, we further analyzed temporal changes of delta power (i.e. density in the delta frequency band during high amplitude low-frequency oscillatory activity, which mirrors sleep depth [25]) in NREM sleep in DBS OFF *vs*. ON conditions for both targets. This analysis revealed a significant average increase of 36% in the average delta power in NREM sleep per 24 h (36.4%, P < 0.05, Wilcoxon’s signed rank tests and Bonferroni corrections; **Fig 5A**) upon VLPO 125 Hz DBS on the ON condition, as compared to DBS OFF. On the other hand, PeFLH 125 Hz DBS induced a significant average decrease of 30% in the average delta power in NREM sleep per 24 h (30.4%, P < 0.05, Wilcoxon’s signed rank tests and Bonferroni corrections; **Fig 5B**).

**Fig 5.**
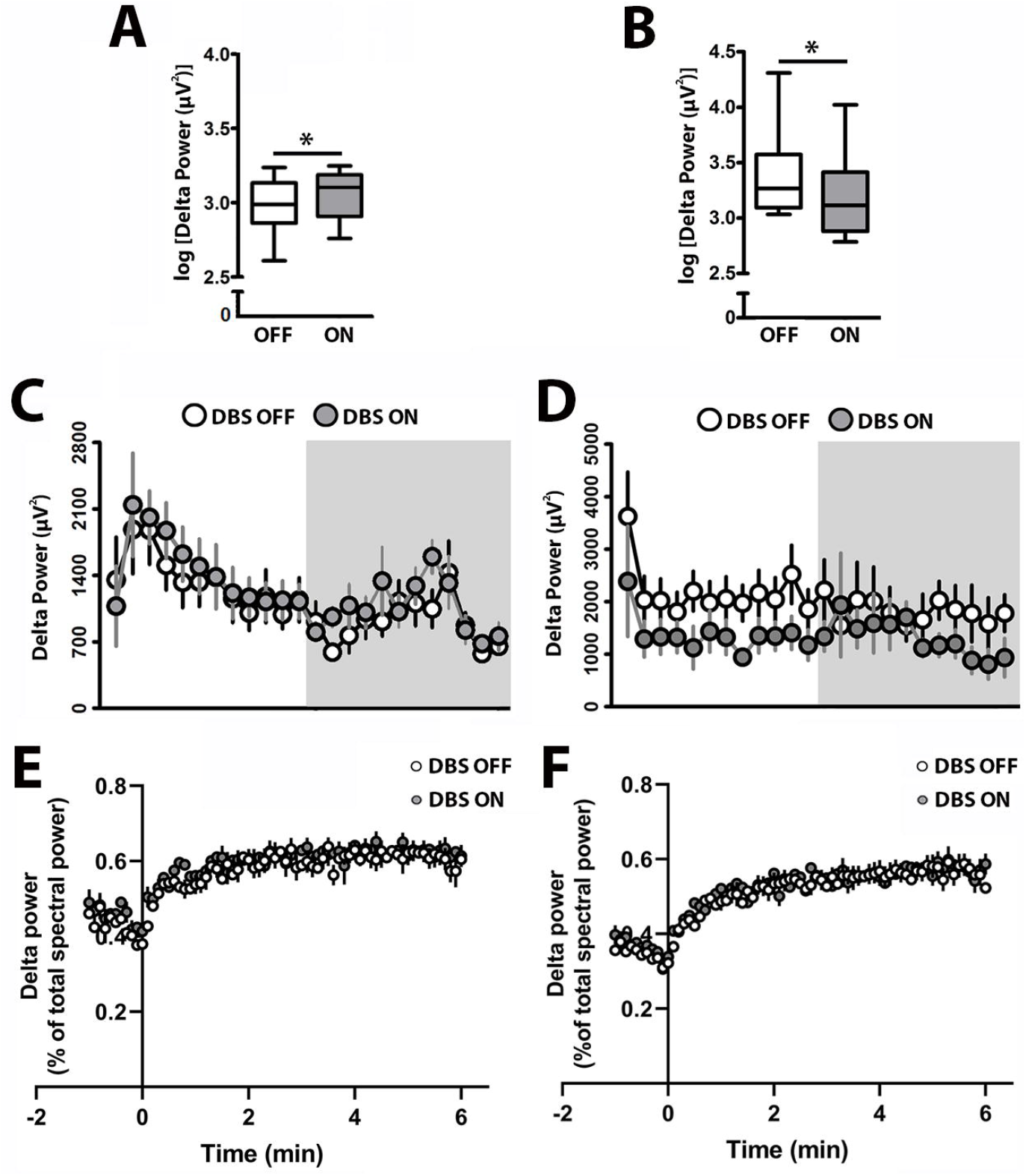
Effect of VLPO and PeFLH HFS modulation on delta power and delta power dynamics during NREM sleep. (**A**) Delta power increased during the 24 h period of recording upon stimulation of VLPO at 125 Hz and decreased (**B**) upon 125 Hz PeFLH stimulation as compared to DBS OFF condition. (**C-D**) The time course of delta power presented in 1 h intervals for 24 h showed no significant effects of DBS at 125 Hz within either VLPO (**C**) or PeFLH (**D**) at any time windows, nor during the light or the dark periods as a whole. (**E**) Neither VLPO nor (**F**) PeFLH DBS affected the build-up of delta power in consolidated, ≥ 6 minutes, NREM episodes. The logarithm of delta power is expressed as medians and quartiles with 95% CI. Wilcoxon’s signed rank tests and Bonferroni corrections. Data in 1 h intervals are means ± S.E.M. * P < 0.05 DBS ON *vs*. DBS OFF. One-way repeated measures ANOVA followed by Student-Newman-Keuls tests. Delta power is reported as absolute values. Two-way repeated measures ANOVA followed by Bonferroni tests. n=6-7 per group. Two statistical outliers (criteria: scores > 2 standard deviations; 1 for the analysis of the light period and 1 for the analysis of the dark period) were excluded from the analysis. Min: minutes; VLPO: ventrolateral preoptic area; PeFLH: perifornical lateral hypothalamic area.

To get more insights into the 24 h time course of delta power changes at HFS, we analyzed the measure in 1 h intervals during both light and dark periods. No specific time windows during the light or dark periods of neither VLPO (**Fig 5C**) nor PeFLH (**Fig 5D**) DBS effects were detected (one-way repeated measures ANOVAs followed by non-significant Student-Newman-Keuls post-hoc comparisons as appropriate; P > 0.05).

To further explore the effect of VLPO and PeFLH HFS on sleep intensity dynamics [23], we calculated the build-up of delta power during consolidated NREM episodes (≥ 6 minutes; **Fig 5E, F**). We observed no significant effect from neither VLPO nor PeFLH stimulation on delta power build-up over consolidated sleep (two-way repeated measures ANOVA, P > 0.05), indicating that changes in delta power were global and not related to its dynamics.

We further explored alterations in the 24 h average full EEG spectrum upon HFS in both target regions, for which we calculated EEG power spectra OFF *vs*. ON stimulation for all 3 vigilance states (**Fig 6**). Despite some variability observed in the EEG spectrum, notably in the delta activity of WAKE, multiple bivariate comparisons for all frequency bands (delta, theta, alpha, beta/gamma) did not reveal significant band-specific changes upon HFS (two-way ANOVA with repeated measures followed by Bonferroni post-hoc comparisons as appropriate, P > 0.05).

**Fig 6.**
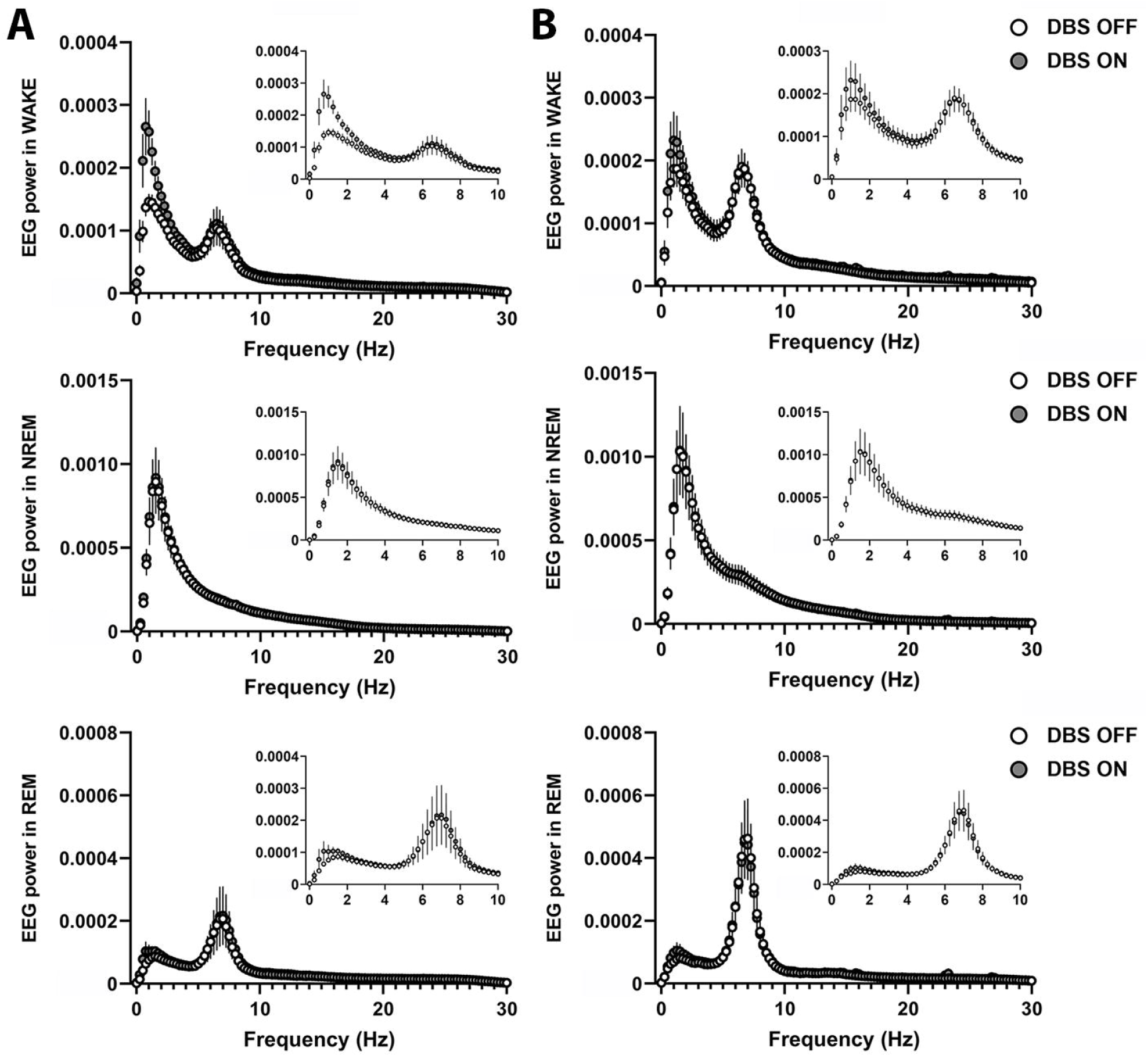
Effect of VLPO and PeFLH DBS on 24 h spectral power in each behavioral state. (**A**) 24 h average spectral power in the EEG during WAKE (top panels), NREM (middle panels) and REM (bottom panels) states upon high-frequency VLPO, or (**B**) PeFLH stimulation. No band-specific effect between DBS and any frequency band were noted. Two-way ANOVA with repeated measures followed by Bonferroni post-hoc comparisons. WAKE: wakefulness, NREM: non-rapid eye movement sleep, REM: rapid eye movement sleep; Hz: hertz; VLPO: ventrolateral preoptic area; PeFLH: perifornical lateral hypothalamic area.

Overall, HFS of VLPO increased delta power, whereas HFS of PeFLH decreased delta power over light and dark periods of recordings.

## Discussion

In this report, we investigated the direct effects over sleep-wake behavior and characteristics of high frequency DBS on sleep- and wake-controlling centers in healthy animals. Our findings suggest that 24 h DBS of the VLPO at 125 Hz modulates sleep-wake characteristics mainly by deepening slow-wave sleep (SWS) as measured by an increase in delta power (36%), while sleep architecture and fragmentation remain unaffected. In other words, the intensity but not the amount of sleep or its stability is enhanced. Similarly, the proportion of vigilance states remained unchanged during 24 h DBS of the wake-promoting PeFLH region but, in contrast to VLPO, stimulation at 125 Hz depressed delta power (30%), weakening SWS. Furthermore, sleep microstructure, as measured by delta build-up over time, was unchanged upon DBS in both targets. Moreover, based on the marked, yet non-significant effect on delta activity upon VLPO DBS, we cannot rule out a behavioral state-specific effect on SWA selectively in wakefulness. This would indicate that VLPO DBS might have a sleep-inducing rather than a sleep-enhancing effect, by acting more on the sleep-inducing permissive neurons rather than on the sleep-maintenance executive neurons [26]. However, due to the small effect and sample size these conclusions remain speculative. The negative results of the detailed spectral power analysis in each behavioral state and the build-up analysis of delta power during consolidated SWS (or NREM) episodes, demonstrated that DBS-elicited changes were global in nature, not affecting sleep intensity dynamics.

### Electrical neuromodulation of VLPO and PeFLH areas

The VLPO has been reported to present a low frequency intrinsic firing pattern of ~10 Hz [27]. Thus, finding increased delta power during NREM sleep with HFS in the VLPO was rather unexpected, given that HFS is empirically known to produce a functional inhibition of the target nucleus. However, this initial oversimplification has been actively disputed, and HFS appears to rely on more complex and multifactorial mechanisms [15, 28, 29]. DBS may notably reduce cellular activity while concurrently exciting the axons of the stimulated neurons, which was shown in both humans and animals [30–32]. Interestingly, excitatory stimulation (i.e. increased output) in parts of the basal ganglia in non-human primates at HFS has shown to excite both glutamatergic [33] and GABAergic [34] efferent neurons. Therefore, it is conceivable that the cell bodies of the VLPO neurons were inhibited by DBS while the output from this GABAergic nucleus increased simultaneously, inhibiting the downstream arousal systems.

VLPO sleep-active neurons have also been shown to progressively increase their firing rate along with sleep depth in rats [35]. The increased intensity of SWS of about 36% upon HFS revealed in our experiments may thus rely on this neurophysiological property. Indeed, HFS could further increase the discharge rate of VLPO sleep-active neurons, based on the synchronization of the target neuronal firing to the stimulus frequency as shown in other nuclei [33], and consequently increase the depth of sleep, as observed in this study.

PeFLH HFS, on the other hand, decreased delta power during NREM sleep by approximately 30%, suggesting an excitatory effect of HFS on this wake-promoting region. Once more, the apparent activation of the PeFLH region at, supposedly inhibitory, HFS is unexpected. Nevertheless, activation of the tuberomammillary nucleus (TMN), a neighboring wake-promoting target with similar firing rates to PeFLH arousal-related neurons, upon HFS (100 Hz) has been already demonstrated in rodents [36]. Moreover, the PeFLH is crossed by the mfbb, carrying projections from groups of neurons critical for sleep-wake control [37], which unintended stimulation could additionally compound in the observed effects. Although the exact neurophysiological mechanisms sustaining these results remain speculative, comparable processes (i.e. activation of neuronal processes rather than cell soma) in VLPO and PeFLH regions may have been involved, increasing the output of these simplistically regarded sleep- and wake-promoting regions by excitation of efferent neurons.

Noteworthily, however, a decrease in SWA has been previously observed by electrical stimulation of the PeFLH neurons [8]. The delivery of trains of electrical stimuli at 50 Hz in the PeFLH area of anesthetized rats increased the mean firing rate of LC neurons and induced an activation of the EEG shown by a decrease in the proportion of delta waves together with an increased percentage of faster (> 4 Hz) waves. This result, although obtained under anesthesia as opposed to our awake, freely-moving rats, reveals a direct role of the PeFLH neurons in the modulation of delta power, in line with our findings.

### Lack of behavioral modulation with VLPO and PeFLH DBS: potential compensatory mechanisms

Although VLPO is an important sleep-promoting center and PeFLH an organizer of wakefulness/sleep stages, the electrical modulation of these regions did not change the proportions of sleep-wake stages, nor the fragmentation of behavioral states in response to HFS. In this line, we cannot exclude a possible DBS effect on regions neighboring VLPO, predominantly populated by wake-active cells in the lateral preoptic / anterior hypothalamic area [35], and/or on the main monoaminergic arousal projections from LC, raphe nuclei and TMN to VLPO, which could counteract the effect of the stimulation on VLPO sleep-active neurons.

Similarly, we cannot exclude the activation of the main inhibitory afferents (i.e. VLPO and median preoptic nucleus) to the PeFLH region upon DBS that could have counterbalanced the stimulatory effects of this conceptualized wake-promoting area. Also, stimulation of a sub-population within the heterogeneous PeFLH, the melanin-concentrating hormone (MCH) neurons, known to promote sleep [38], might have played a role. Indeed, these neurons - electrically silent in the absence of synaptic activity - show pronounced firing at 100 Hz upon brief repeated current injections in rodent brain slices [39, 40]. Strikingly, application of high levels of orexin peptides excited some MCH neurons in vitro [41]. In this context, possible over-excitation of orexin neurons by electrical stimulation may have in turn stimulated MCH neurons as a feedback mechanism to prevent hyperarousal and, consequently, overall changes in sleep-wake amounts.

### Limitations

The most challenging part of any DBS study is identifying the optimal combination of parameters producing a targeted effect while limiting side effects, for which the lack of conclusive dose-response assessments for each parameter analyzed (frequency, intensity, duration) is one of the main limitations of our study. Also, despite the use of bipolar electrodes producing a concentrated current around the tip of the electrodes, our study expectedly suffers from a fundamental DBS limitation which, unlike other highly specific approaches such as optogenetics [42], lacks selectivity between the activation of local cells and that of passing axons using conventional symmetrical biphasic pulses, as illustrated in computational models with monopolar electrodes [43]. Additionally, cell heterogeneity within a target constitutes an intrinsic biological limitation of DBS. For instance, electrical stimulation of PeFLH, whose activation has been shown to produce changes at feeding and motor activity [44], could have additionally affected another sub-population besides the wake-promoting hypocretin neurons, such as MCH neurons, known to discharge in a reciprocal manner to orexin neurons across the sleep-wake cycle [45, 46] and promote sleep [38]. Nevertheless, DBS remains to be a preferred tool of clinical choice, as opposed to approaches involving invasive mutagenic and viral strategies. For this, its exploration remains remarkably important in the context of the search for new therapeutic targets.

A limitation of technical nature is the use of a relatively short pulse width in our study. Longer pulse widths (in the range of milliseconds) could be used in the future to preferentially stimulate the cell bodies of our targets [47]. However, increasing the pulse width would also increase the charge density (i.e. the amount of electrical charge per surface area) accumulated per pulse, thus potentially reaching safety limits and causing tissue lesions, which we avoided in our study.

## Conclusions

Our investigation has provided new insights into DBS modulation of important centers of the sleep-wake-regulating network. In summary, our unexpected results suggest that tight compensatory mechanisms counteracted the intended changes in sleep-wake behavior in our healthy rats. However, escaping the tight regulatory controls, sleep intensity is specifically – but contrary to predicted – modulated by hypothalamic DBS, shining light on non-acute neuromodulation outputs of hypothalamic high frequency electrical stimulation. Overall, although our results indicate that clinical implementation of HFS in hypothalamic centers for the treatment of disabling sleep-wake behavior disorders should eventually be explored with utmost caution, they encourage new avenues of research intended to further explore and understand the findings for instance by means of implementation of hypothalamic HFS in animal models of sleep-wake disorders.

## Acknowledgments

We thank Dr. O. Sürücü for contribution to study concept and Mr. T. Provini and Dr. I. Amrein for technical support.

## Research funding

This work was supported by the HSM-II program of the Canton of Zurich (CRB), by the Forschungskredit of the University of Zurich (DN), and by the Clinical Research Priority Program Sleep and Health of the University of Zurich (CRB).

## Conflict of interest

Authors state no conflict of interest.

## Data availability statement

The datasets generated during and/or analyzed during the current study are available from the corresponding author on reasonable request.

## Notes

### Competing Interest Statement

The authors have declared no competing interest.

